# PBxplore: a tool to analyze local protein structure and deformability with protein Blocks

**DOI:** 10.1101/136408

**Authors:** Jonathan Barnoud, Hubert Santuz, Pierrick Craveur, Agnel Praveen Joseph, Vincent Jallu, Alexandre G. de Brevern, Pierre Poulain

## Abstract

Proteins are highly dynamic macromolecules. A classical way to analyze their inner flexibility is to perform molecular dynamics simulations. In this context, we present the advantage to use small structural prototypes, namely the Protein Blocks (PBs). PBs give a good approximation of the local structure of the protein backbone. More importantly, by reducing the conformational complexity of protein structures, they allow analyzes of local protein deformability which cannot be done with other methods and had been used efficiently in different applications. PBxplore is a suite of tools to analyze the dynamics and deformability of protein structures using PBs. It is able to process large amount of data such as those produced by molecular dynamics simulations. It produces various outputs with text and graphics, such as frequencies, entropy and information logo. PBxplore is available at https://github.com/pierrepo/PBxplore and is released under the open-source MIT license.

## INTRODUCTION

Proteins are highly dynamic macromolecules (Frauenfelder et al., 1991; Bu and Callaway, 2011). To analyze their inner flexibility, computational biologists often use molecular dynamics (MD) simulations. The quantification of protein flexibility is based on various methods such as Root Mean Square Fluctuations (RMSF) that relies on multiple MD snapshots or Normal Mode Analysis (NMA) that relies on a single structure and focus on quantifying large movements.

Other interesting *in silico* approaches assess protein motions through the protein residue network (Atilgan et al., 2007) or dynamical correlations from MD simulations (Ghosh and Vishveshwara, 2007; Dixit and Verkhivker, 2011). We can also notice the development of the MOdular NETwork Analysis (MONETA), which localizes the perturbations propagation throughout a protein structure (Laine et al., 2012).

Here we use an alternative yet powerful approach based on small prototypes or “structural alphabets” (SAs). SAs approximate conformations of protein backbones and code the local structures of proteins as one-dimensional sequences (Offmann et al., 2007). Protein Blocks (PBs) (de Brevern et al., 2000) is one of these SAs (de Brevern, 2005; Etchebest et al., 2005; Joseph et al., 2010).

PBs are composed of 16 blocks designed through an unsupervised training performed on a representative non-redundant databank of protein structures (de Brevern et al., 2000). PBs are defined from a set of dihedral angles describing the protein backbone. This property makes PBs interesting conformational prototypes of the local protein structure. PBs are labeled from *a* to *p* (see Fig. 1a). PBs *m* and *d* are prototypes for central α-helix and central *β* -strand, respectively. PBs *a* to *c* primarily represent *β* -strand N-caps and PBs *e* and *f*, *β* -strand C-caps; PBs *g* to *j* are specific to coils, PBs *k* and *l* are specific to α-helix N-caps, and PBs *n* to *p* to α-helix C-caps (de Brevern, 2005). Figure 1 illustrates how a PB sequence is assigned from a protein structure. Starting from the 3D coordinates of the barstar protein (Fig. 1b), the local structure of each amino acid is compared to the 16 PB definitions (Fig. 1a). The most similar protein block is assigned to the residue under consideration (the similarity metric is explained latter in this article). Eventually, assignment leads to the PB sequence represented in Fig. 1c.

**Figure 1:**
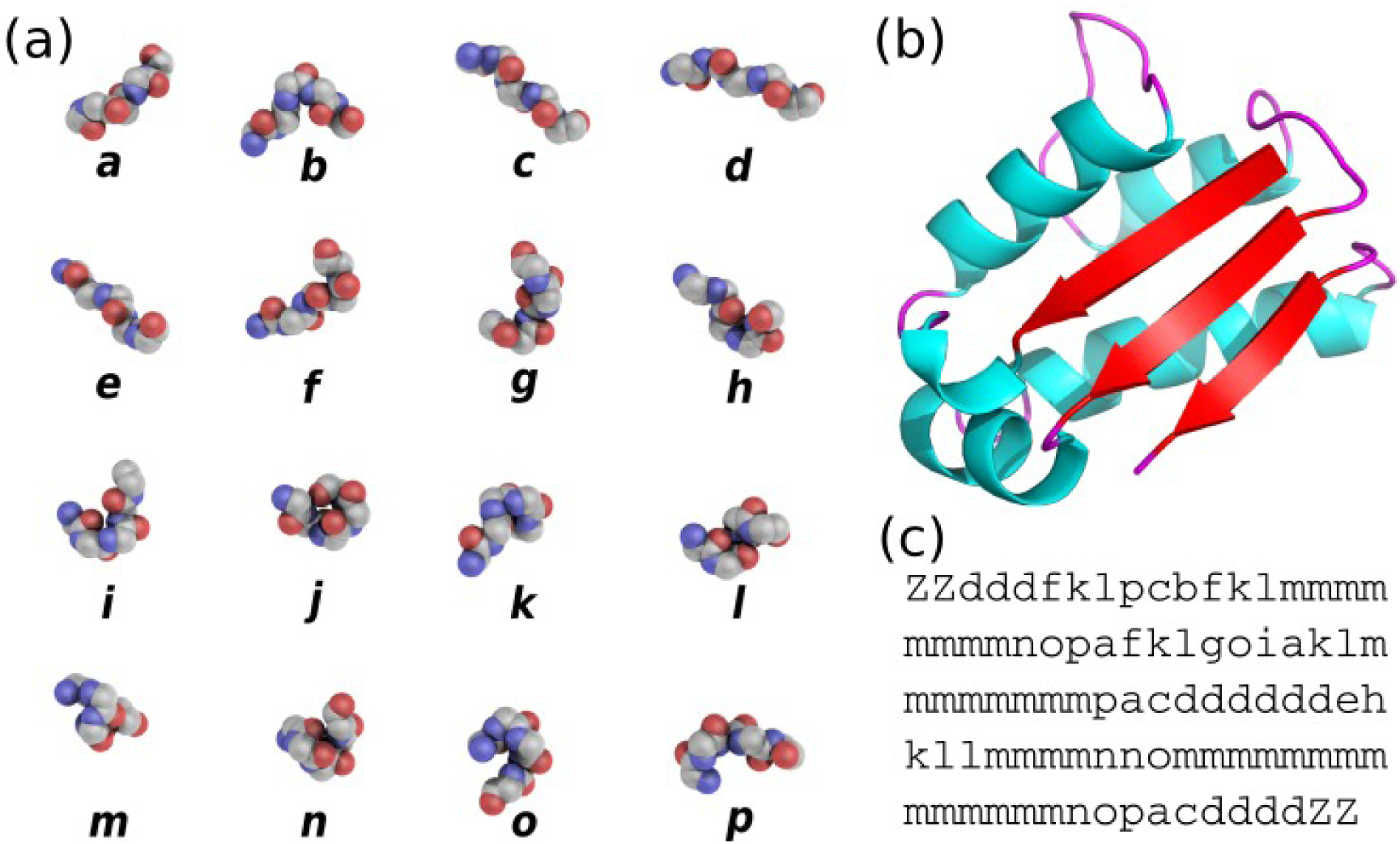
(a) The 16 protein blocks (PBs) represented in balls with carbon atoms in gray, oxygen atoms in red and nitrogen atoms in purple (hydrogen atoms are not represented). (b) The barstar protein (PDB ID 1AY7 (Sevcík et al., 1998)) represented in cartoon with alpha-helices in blue, beta-strands in red and coil in pink. These representations were generated using PyMOL software (DeLano, 2002) (c) PBs sequence obtained from PBs assignment. Z is a dummy PB meaning that no PB can be assigned to this position.

By reducing the complexity of protein structure, PBs have been showed to be efficient and relevant in a wide spectrum of applications. To name a few, PBs have been used to analyze protein contacts (Faure et al., 2008), to propose a structural model of a transmembrane protein (de Brevern, 2005), to reconstruct globular protein structures (Dong et al., 2007), to design peptides (Thomas et al., 2006), to define binding site signatures (Dudev and Lim, 2007), to perform local protein conformation predictions (Li et al., 2009; Rangwala et al., 2009; Suresh et al., 2013; Suresh and Parthasarathy, 2014; Zimmermann and Hansmann, 2008), to predict *β* -turns (Nguyen et al., 2014) and to understand local conformational changes due to mutations of the αIIb*β* 3 human integrin (Jallu et al., 2012, 2013, 2014).

PBs are also useful to compare and superimpose protein structures with pairwise and multiple approaches (Joseph et al., 2011, 2012), namely iPBA (Gelly et al., 2011) and mulPBA (Léonard et al., 2014), both currently showing best results compared to other superimposition methods. Eventually, PBs lead to interesting results at predicting protein structures from their sequences (Ghouzam et al., 2015, 2016) and at predicting protein flexibility (Bornot et al., 2011; de Brevern et al., 2012).

Our results on biological systems such as, the DARC protein (de Brevern et al., 2005), the human αIIb*β* 3 integrin (Jallu et al., 2012, 2013, 2014) and the KISSR1 protein (Chevrier et al., 2013), highlighted the usefulness of PBs to understand local deformations of large protein structures. Specially, these analyzes have shown that a region considered as highly flexible through RMSF quantifications, can be seen through PBs as locally highly rigid. This unexpected behavior is explained by a local rigidity, surrounded by deformable regions (Craveur et al., 2015). To go further, we recently used PBs to analyze long-range allosteric interactions in the Calf-1 domain of αIIb integrin (Goguet et al., 2017). To our knowledge, the only other related approach based on SA to assess local deformation is GSATools (Pandini et al., 2013), it is specialized in the analysis of functional correlations between local and global motions, and the mechanisms of allosteric communication.

Despite the versatility of PBs and the large spectrum of their applications, PBs lack a uniform and easy-to-use toolkit to assign PB sequences from 3D structures, and to analyze these sequences. The only known implementation is a an old C program not publicly available and not maintained anymore. Such tool not being available reduces the availability of the PBs for studies where they would be meaningful.

We thus propose PBxplore, a tool to analyze local protein structure and deformability using PBs. It is available at https://github.com/pierrepo/PBxplore. PBxplore can read PDB structure files (Bernstein et al., 1977), PDBx/mmCIF structure files (Bourne et al., 1997), and MD trajectory formats from most MD engines, including Gromacs MD topology and trajectory files (Lindahl et al., 2001; van der Spoel et al., 2005). Starting from 3D protein structures, PBxplore assigns PBs sequences; computes a local measurement of entropy, a density map of PBs along the protein sequence and a WebLogo-like representation of PBs.

In this paper, we first present the principle of PBxplore, then its different tools, and finally a step-by-step user-case with the *β* 3 subunit of the human platelet integrin αIIb*β* 3.

## DESIGN AND IMPLEMENTATION

PBxplore is written in Python (van Rossum, 1995; Software, 2010; Bassi, 2007). It is compatible with Python 2.7, and with Python 3.4 or greater. It requires the Numpy Python library for array manipulation (Ascher et al., 1999), the matplotlib library for graphical representations, and the MDAnalysis library for molecular dynamics simulation files input (Michaud-Agrawal et al., 2011; Gowers et al., 2016). Optionally, PBxplore functionalities can be enhanced by the installation and the use of WebLogo (Crooks et al., 2004) to create sequence logos.

PBxplore is available as a set of command-line tools and as a Python module. The command-line tools allow for an easy integration of PBxplore in existing analysis pipelines. These programs can be linked up together to carry out the most common analyses on PB sequences to provide insights on protein flexibility. In addition, the PBxplore Python library provides an API to access its core functionalities which allows the integration of PBxplore in Python programs and workflows, and the extension of the method to suit new needs.

PBxplore is released under the open-source MIT license (Open Source Initiative, 2014). It is available on the software development platform GitHub (GitHub, 2007) at https://github.com/pierrepo/PBxplore.

The package contains unit and regression tests and is continuously tested using Travis CI (Travis CI, 2015). An extensive documentation is available on Read the Docs (Holscher et al., 2010) at https://pbxplore.readthedocs.io.

### Installation

The easiest way to install PBxplore is through the Python Package Index (PyPI):

~~~
pip install --user pbxplore
~~~

It will ensure all required dependencies are installed correctly.

### Command-line Tools

A schematic description of PBxplore command line interface is provided in Fig. 2. The interface is composed of three different programs: `PBassign` to assign PBs, `PBcount` to compute PBs frequency on multiple conformations, and `PBstat` to perform statistical analyses and visualization. These programs can be linked up together to make a structure analysis pipeline to study protein flexibility.

**Figure 2:**
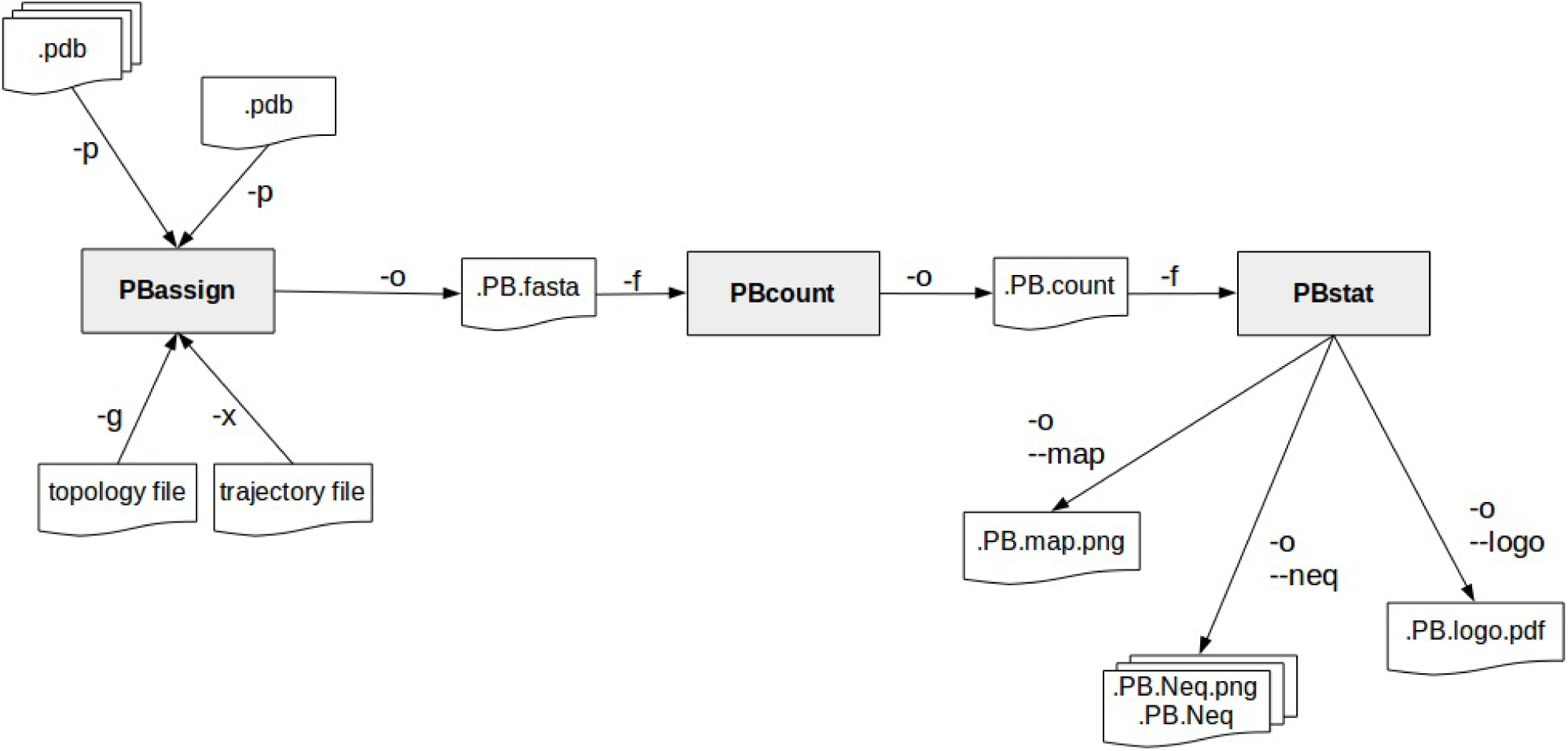
PBxplore is based on 3 programs that can be chained to build a structure analysis pipeline. Main input file types (.pdb, MD trajectory, MD topology), output files (.fasta,.png,.Neq,.pdf) and parameters (beginning with a single or double dash) are indicated.

### PBassign

The very first task is to assign PBs from the protein structure(s). A PB is associated to each pentapeptide included in the protein sequence. To assign a PB to a residue *n*, 5 residues are required (residues *n* − 2, *n* − 1, *n*, *n* + 1 and *n* + 2). From the structure of these 5 residues, 8 dihedral angles (ψ and ϕ) are computed, going from the ψ angle of residue *n* − 2 to the ϕ angle of residue *n* + 2 (de Brevern, 2005). This set of 8 dihedral angles is then compared to the reference angles set of the 16 PBs (de Brevern et al., 2000) using the Root Mean Square Deviation Angle (RMSDA) measure, i.e., an Euclidean distance on angles. PB with the smallest RMSDA is assigned to residue *n*. A dummy PB *Z* is assigned to residues for which all 8 angles cannot be computed. Hence, the first two N-terminal and the last two C-terminal residues are always assigned to PB *Z*.

The program `PBassign` reads one or several protein 3D structures and performs PBs assignment as one PBs sequence per input structure. `PBassign` can process multiple structures at once, either provided as individual structure files, as a directory containing many structure files or as topology and trajectory files issued from MD simulations. Note that PBxplore is able to read any trajectory file format handled by the MDAnalysis library, yet our tests focused on Gromacs trajectories. Output PBs sequences are bundled in a single file in fasta format.

### PBcount

During the course of a MD simulation, the local protein conformations can change. It is then interesting to analyze them through PB description. Indeed, as each PB describes a local conformation, the variability of the PB assigned to a given residue throughout the trajectory indicates some local deformation of the protein structure. Thus, once PBs are assigned, PBs frequencies per residue can be computed.

The program `PBcount` reads PBs sequences for different conformations of the same protein from a file in the fasta format (as outputted by `PBassign`). Many input files can be provided at once. The output data is a 2D matrix of *x* rows by *y* columns, where *x* is the length of the protein sequence and *y* is the 16 distinct PBs. A matrix element is the count of a given PB at a given position in the protein sequence.

### PBstat

The number of possible conformational states covered by PBs is higher than the classical secondary structure description (16 states instead of 3). As a consequence, the amount of information produced by `PBcount` can be complex to handle. Hence, we propose three simple ways to visualize the variation of PBs which occur during a MD simulation.

The program PBstat reads PBs frequencies as computed by PBcount. It can produce three types of outputs based on the input argument(s). The first two use the matplotlib library and the last one requires the installation of the third-party tool Weblogo (Crooks et al., 2004). PBstat also offers two options (`--residue-min` and `--residue-max`) to define a residue frame allowing the user to quickly look at segments of interest. The three graphical representations proposed are:

- *Distribution of PBs.* This feature plots the frequency of each PB along the protein sequence. The output file could be in format.png,.jpg or.pdf. A dedicated colorblind safe color range (Brewer et al., 2013) allows visualizing the distribution of PBs. For a given position in the protein sequence, blue corresponds to a null frequency when the particular PB is never sampled at this position and red corresponds to a frequency of 1 when the particular PB is always found at this position. This representation is produced with the `--map` argument.
- *Equivalent number of PBs (N*_*eq*_). The *N*_*eq*_ is a statistical measurement similar to entropy (Offmann et al., 2007). It represents the average number of PBs sampled by a given residue. *N*_*eq*_ is calculated as follows:

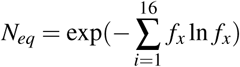

where *f*_*x*_ is the probability (or frequency) of the PB *x*. A *N*_*eq*_ value of 1 indicates that only a single type of PB is observed, while a value of 16 is equivalent to a random distribution, i.e. all PBs are observed with the same frequency 1/16. For example, a *N*_*eq*_ value around 5 means that, across all the PBs observed at the position of interest, 5 different PBs are mainly observed. If the *N*_*eq*_ exactly equals to 5, this means that 5 different PBs are observed in equal proportions (i.e. 1/5).
A high *N*_*eq*_ value can be associated with a local deformability of the structure whereas a *N*_*eq*_ value close to 1 means a rigid structure. In the context of structures issued from MD simulations, the concept of deformability / rigidity is independent to the one of mobility. The *N*_*eq*_ representation is produced with the `--neq` argument.
- *Logo representation of PBs frequency.* This is a WebLogo-like representation (Crooks et al., 2004) of PBs sequences. The size of each PB is proportional to its frequency at a given position in the sequence. This type of representation is useful to pinpoint PBs patterns. This WebLogo-like representation is produced with the `--logo` argument.

### Python Module

PBxplore is also a Python module that more advanced users can embed in their own Python script. Here is a Python 3 example that assigns PBs from the structure of the barstar ribonuclease inhibitor (Lubienski et al., 1994):

~~~
import urllib.request
import pbxplore as pbx
# Download the pdb file urllib.request.urlretrieve(’https://files.rcsb.org/view/1BTA.pdb’, ’1BTA.pdb’)
# The function pbx.chain_from_files() reads a list of files
# and for each one returns the chain and its name.
for chain_name, chain in pbx.chains_from_files([’1BTA.pdb’]):
# Compute phi and psi angles
dihedrals = chain.get_phi_psi_angles()
# Assign PBss
pb_seq = pbx.assign(dihedrals)
print(’PBs sequence for chain {}:\n{}’.format(chain_name, pb_seq))
~~~

The documentation contains complete and executable Jupyter notebooks explaining how to use the module. It goes from the PBs assignments to the visualization of the protein deformability using the analysis functions. This allows the user to quickly understand the architecture of the module.

## RESULTS

This section aims at giving the reader a quick tour of PBxplore features on a real-life example. We will focus on the *β* 3 subunit of the human platelet integrin αIIb*β* 3 that plays a central role in hemostasis and thrombosis. The *β* 3 subunit has also been reported in cases of alloimmune thrombocytopenia (Kaplan, 2006; Kaplan and Freedman, 2007). We studied this protein by MD simulations (for more details, see references (Jallu et al., 2012, 2013, 2014)).

The *β* 3 integrin subunit structure (Poulain and de Brevern, 2012) comes from the structure of the integrin complex (PDB 3FCS (Zhu et al., 2008)). Final structure has 690 residues and was used for MD simulations. All files mentioned below are available in the `demo_paper` directory from the GitHub repository (https://github.com/pierrepo/PBxplore/tree/master/demo_paper).

### Protein Blocks assignment

The initial file `beta3.pdb` contains 225 structures issued from a single 50 ns MD simulation of the *β* 3 integrin.

~~~
PBassign -p beta3.pdb -o beta3
~~~

This instruction generates the file `beta3.PB.fasta`. It contains as many PB sequences as there are structures in the input `beta3.pdb` file.

Protein Blocks assignment is the slowest step. In this example, it took roughly 80 seconds on a laptop with a quad-core-1.6-GHz processor.

### Protein Blocks frequency

~~~
PBcount -f beta3.PB.fasta -o beta3
~~~

The above command line produces the file `beta3.PB.count` that contains a 2D-matrix with 16 columns (as many as different PBs) and 690 rows (one per residue) plus one supplementary column for residue number and one supplementary row for PBs labels.

### Statistical analysis

#### Distribution of PBs

~~~
PBstat -f beta3.PB.count -o beta3 --map
~~~

Figure 3 shows the distribution of PBs for the *β* 3 integrin. The color scale ranges from blue (the PB is not found at this position) to red (the PB is always found at this position). The *β* 3 protein counts 690 residues. This leads to a cluttered figure and prevents getting any details on a specific residue (Fig. 3a). However, it exhibits some interesting patterns colored in red that correspond to series of neighboring residues exhibiting a fixed PB during the entire MD simulation. See for instance patterns associated to PBs *d* and *m* that reveal *β* -sheets and α-helices secondary structures (de Brevern, 2005).

**Figure 3:**
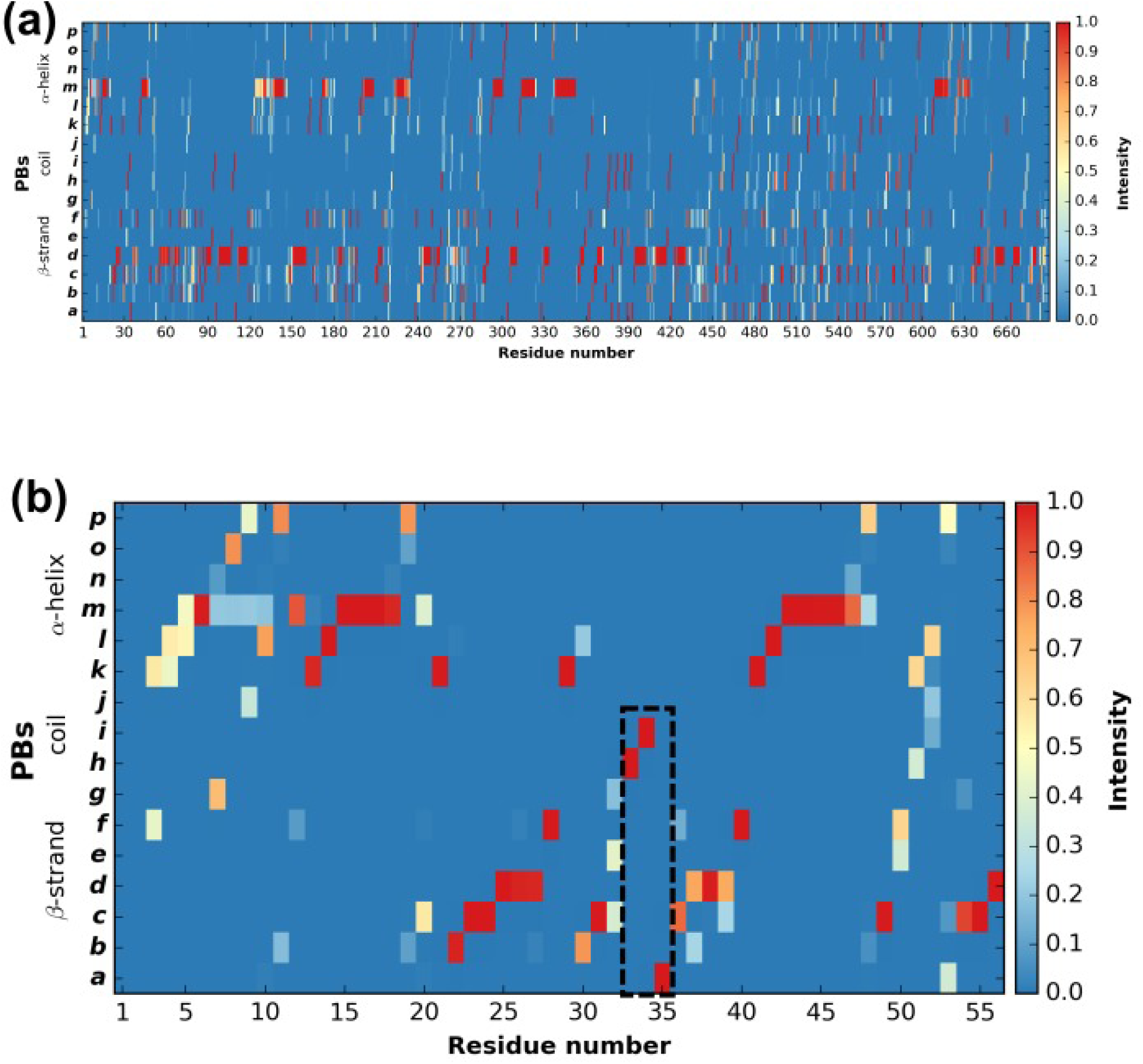
Distribution of PBs for the *β* 3 integrin along the protein sequence. On the x-axis are found the 690 position residues and on the y-axis the 16 consecutive PBs from *a* to *p* (the two first and two last positions associated to “Z” have no assignment). (a) For the entire protein. (b) For the PSI domain only (residues 1 to 56). The dashed zone pinpoints residue 33 to 35.

With a large protein such as this one, it is better to look at limited segments. A focus on the PSI domain (residue 1 to 56) (Jallu et al., 2012; Zhu et al., 2008) of the *β* 3 integrin was achieved with the command:

~~~
PBstat -f beta3.PB.count -o beta3 --map --residue-min 1 --residue-max 56
~~~

Figure 3b shows the PSI domain dynamics in terms of PBs. Interestingly, residue 33 is the site of the human platelet antigen (HPA)-1 alloimmune system. It is the first cause of alloimmune thrombocytopenia in Caucasian populations and a risk factor for thrombosis (Kaplan, 2006; Kaplan and Freedman, 2007). In Fig. 3b, this residue occupies a stable conformation with PB *h*. Residues 33 to 35 define a stable core composed of PBs *h-i-a*. This core is found in all of the 255 conformations extracted from the MD simulation and then is considered as highly rigid. On the opposite, residue 52 is flexible as it is found associated to PBs *i*, *j*, *k* and *l* corresponding to coil and α-helix conformations.

#### Equivalent number of PBs

The *N*_*eq*_ is a statistical measurement similar to entropy and is related to the flexibility of a given residue. The higher is the value, the more flexible is the backbone. The *N*_*eq*_ for the PSI domain (residue 1 to 56) was obtained from the command line:

~~~
PBstat -f beta3.PB.count -o beta3 --neq --residue-min 1 --residue-max 56
~~~

The output file `beta3.PB.Neq.1-56` contains two columns, corresponding to the residue numbers and the *N*_*eq*_ values. Figure 4a represents the *N*_*eq*_ along with the PBs sequence of the PSI domain, as generated by `PBstat`. The rigid region 33-35 and the flexible residue 52 are easily spotted, with low *N*_*eq*_ values for the former and a high *N*_*eq*_ value for the latter.

**Figure 4:**
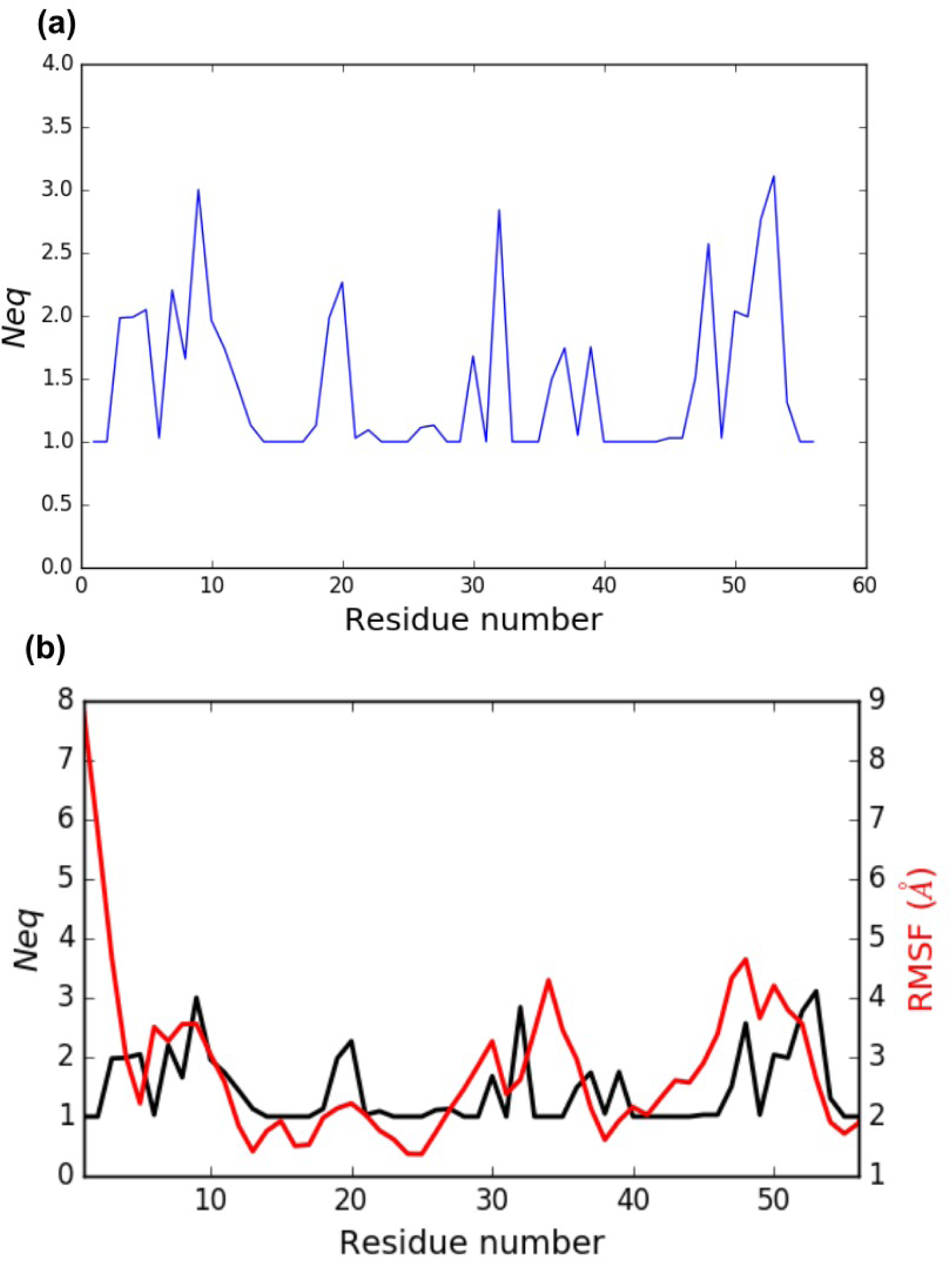
(a) *N*_*eq*_ versus residue number for the PSI domain (residues 1 to 56). (b) Comparison between RMSF and *N*_*eq*_.

An interesting point, seen in our previous studies, is that the region delimited by residues 33 to 35 was shown to be highly mobile by the RMSF analysis we performed in Jallu et al. (2012) (for more details, see Materials and Methods section in Jallu et al. (2012)). For comparison, RMSF and *N*_*eq*_ are represented on the same graph on Fig. 4b. This high mobility was correlated with the location of this region in a loop, which globally moved a lot in our MD simulations. Here, we observe that the region 33-35 is rigid. The high values of RMSF we observed in our previous work were due to flexible residues in the vicinity of the region 33-35, probably acting as hinges (residues 32 and 36–37). Understanding the flexibility of residues 33 to 35 is important since this region defines the HPA-1 alloantigenic system involved in severe cases of alloimmune thrombocytopenia. PBxplore allows discriminating between flexible and rigid residues. The *N*_*eq*_ is a metric of deformability and flexibility whereas RMSF quantifies mobility.

#### Logo representation of PBs frequency

While the *N*_*eq*_ analysis focuses on the flexibility of amino acids, the WebLogo-like representation (Crooks et al., 2004) aims at identifying the diversity of PBs and their frequencies at a given position in the protein sequence. With a focus on the PSI domain, the following command line was used:

~~~
PBstat -f beta3.PB.count -o beta3 --logo --residue-min 1 --residue-max 56
~~~

Figure 5 represents PBs found at a given position. The rigid region 33-35 is composed of a succession of PBs *h-i-a* while the flexible residue 52 is associated to PBs *i*, *j*, *k* and *l*. This third representation summarized pertinent information, as shown in Jallu et al. (2013).

**Figure 5:**
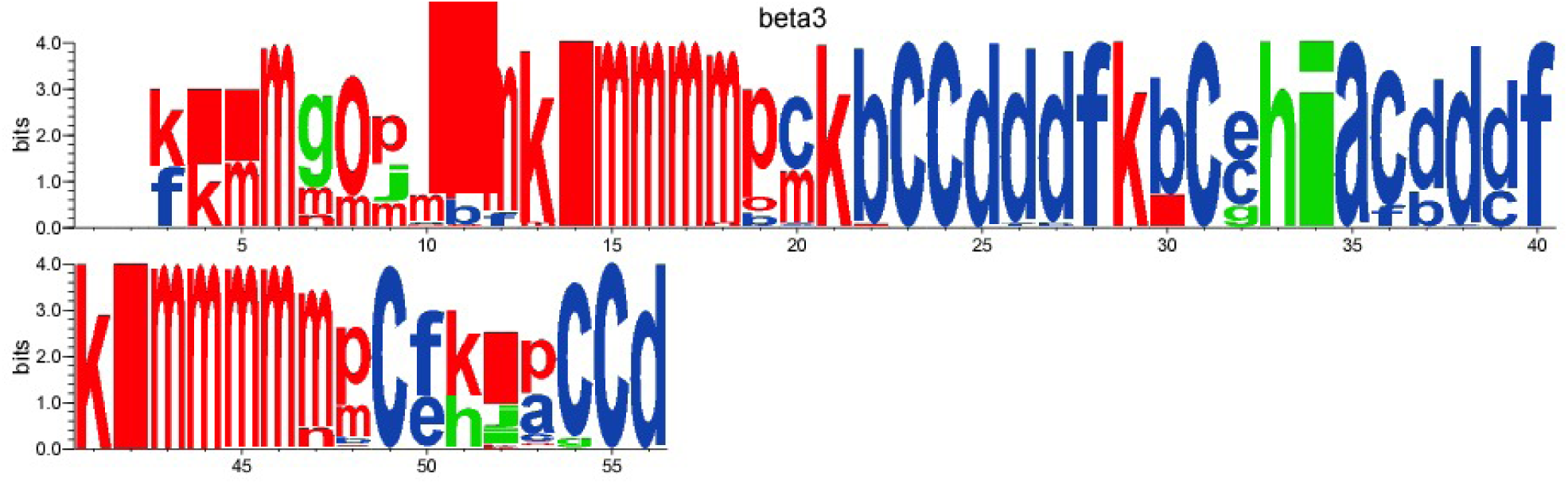
WebLogo-like representation of PBs for the PSI domain of the *β* 3 integrin. PBs in red roughly correspond to α-helices, PBs in blue to *β* -sheets and PBs in green to coil.

## CONCLUSION

From our previous works (Jallu et al., 2012, 2013, 2014; Chevrier et al., 2013), we have seen the usefulness of a tool dedicated to the analysis of local protein structures and deformability with PBs. We also showed the relevance of studying molecular deformability in the scope of structures issued from MD simulations. In a very recent study (Goguet et al, 2017), long independent MD simulations were performed for seven variants and one reference structure of the Calf-1 domain of the αIIb human integrin. Simulations were analyzed with PBxplore. Common and flexible regions as well as deformable zones were observed in all the structures. The highest B-factor region of Calf-1, usually considered as most flexible, is in fact a rather rigid region encompassed into two deformable zones. Each mutated structure barely showed any modifications at the mutation sites while distant conformational changes were detected by PBxplore. These results question the relationship between MD simulations and allostery and the role of long range effects on protein structure. In this context, we propose PBxplore, freely available at https://github.com/pierrepo/PBxplore. It is written in a modular fashion that allows embedding in any PBs related Python application.

### Software Availability

PBxplore is released under the open-source MIT license (Open Source Initiative, 2014). Its source code can be freely downloaded from the GitHub repository of the project: https://github.com/pierrepo/PBxplore. In addition, the present version of PBxplore (1.3.7) is also archived in the digital repository Zenodo (Barnoud et al., 2017).

## ACKNOWLEDGEMENTS

This work was supported by grants from National Institute for Blood Transfusion (INTS, France) and Lab of Excellence GR-Ex to JB, PC, SL, APJ, VJ, AGdB and PP, from the Ministry of Research (France), University Paris Diderot, Sorbonne Paris Cité(France), National Institute for Health and Medical Research (INSERM, France) to JB, PC, SL, APJ, AGdB and PP. The labex GR-Ex, reference ANR-11-LABX-0051 is funded by the program ”Investissements d’avenir” of the French National Research Agency, reference ANR-11-IDEX-0005-02. AGdB acknowledges to Indo-French Centre for the Promotion of Advanced Research / CEFIPRA for collaborative grant (number 5302-2). JB was supported by the TOP program of Prof. Marrink, financed by the Netherlands Organisation for Scientific Research (NWO).

## AUTHOR CONTRIBUTIONS

PP and AGdB conceived the project. PP, JB and HS wrote the software. AGdB, PC, APJ and VJ improved and tested the software. All authors reviewed the manuscript.

